# EPIGENETIC REPROGRAMMING OF CELL IDENTITY IN THE RAT PRIMARY NEURON-GLIA CULTURES INVOLVES HISTONE SEROTONYLATION

**DOI:** 10.1101/2025.05.13.653697

**Authors:** A.A. Borodinova, Yu.A. Leontovich, A.P. Beletskiy, A.V. Revishchin, G.V. Pavlova, P.M. Balaban

## Abstract

Epigenetic rearrangements can create a favorable environment for the intrinsic plasticity of brain cells, leading to cellular reprogramming into virtually any cell type through the induction of cell-specific transcriptional programs. In this study, we assessed how chromatin remodeling induced by broad-spectrum HDAC inhibitors affects cellular differentiation trajectories in rat primary neuron-glia cultures using a combination of transcriptomics, qPCR and cytochemistry. We described epigenetic regulation of transcriptional programs controlled by master transcription factors and neurotrophins in the context of neuronal and glial differentiation and evaluated the expression of representative cell-specific markers. The results obtained suggest that HDAC inhibitors reduce proliferative potential of cultured cells and induce transcriptomic changes associated with cell differentiation and specialization. Particularly, we revealed a significant upregulation of genes typically expressed in neuromodulatory neurons, and downregulation of genes expressed in glia and inhibitory neurons. Transcriptional changes were accompanied by continuous elevation of histone serotonylation levels in both neurons and glia. We assume that early appearance of the enhanced histone serotonylation marks, and the persistence of these changes over many hours in distinct brain cells may indicate that chromatin remodeling induced by histone serotonylation contributes to the maintainance of a new transcriptional programs associated with cellular reprogramming.

## Introduction

The multistep differentiation of neural progenitor cells into specific cell types (neurons, astrocytes, oligodendrocytes) in the developing brain requires epigenetic rearrangements of chromatin landscape capable of altering specific gene expression programs (Juliandi et al., 2010). The level of histone acetylation is particularly important for cell lineage specification. Developing glial cells (oligodendrocytes, astrocytes) show the reduced histone acetylation levels, as compared to neural progenitors and immature neurons (Hsieh et al., 2004).

It is well established that epigenetic regulators from the histone deacetylase (HDAC) family govern cells’ differentiation throughout brain development and later in adulthood. Particularly, active HDACs play an important role in transcriptional inactivation of neuron-specific genes in nonneuronal cells (Huang et al., 1999). Numerous studies have examined the role of specific HDACs in regulation of glial and neuronal fates in the course of cell differentiation (Humphrey et al., 2008; Ye et al., 2009; Conway et al., 2012; Castelo-Branco et al., 2014; Zhang et al., 2016). At early developmental stages corresponding to high rate of differentiation of multipotent neural stem cells (NSCs), members of class I HDACs (HDAC1-3) appear to act redundantly to negatively regulate oligodendrocyte differentiation, which is rescued by inactivation of either HDAC1, HDAC2 or HDAC3 (Humphrey et al., 2008; Castelo-Branco et al., 2014). It has been proposed that HDAC1 transcriptionally controls the genes responsible for neurogenic programs (Harrison et al., 2011), and its deletion affects neuronal differentiation of multipotent NSCs (Nieto-Estevez et al., 2022). In contrast, HDAC3 removal initiated neuronal differentiation of multipotent NSCs (Castelo-Branco et al., 2014), while HDAC3 overexpression selectively stimulated differentiation of multipotent NSCs to astrocytes (Humphrey et al., 2008). At a later stage of development, the regulatory role of HDACs is reversed. Thus, HDACs class I restricted the lineage commitment of the oligodendrocyte precursor cells (OPCs), and induced the oligodendrocyte differentiation both *in vitro* and *in vivo* (Ye et al., 2009; Zhang et al., 2016). Moreover, both HDAC1 and HDAC2 were necessary for neuronal differentiation of neural progenitor cells *in vivo* (NPCs) (Montgomery et al., 2009). It has been noticed that HDAC3 acts bi-directionally as a molecular switch for oligodendrocytes and astrocytes fate decision: its cooperation with p300 histone acetyltransferase stimulates oligodendrocytes differentiation, while selective blockade of HDAC3 facilitates astrocytes lineage specification in OPCs (Zhang et al., 2016). Therefore, these data imply a changing role of HDACs during brain cell differentiation, which requires a specific time window for proper regulation of cell identity.

Considerable efforts were done to investigate the role of epigenetics in cell differentiation using the broad-spectrum HDAC inhibitors. It was found that HDAC inhibitors can trigger developmental plasticity in primary OPC cultures: their application prevents oligodendrocyte differentiation and astrocyte fate commitment (Conway et al., 2012), as well as activates proneural genes that revert OPCs into multipotent neural stem cells (Lyssiotis et al., 2007). This effect may probably induce subsequent reprogramming into other cell types, but it requires further investigation. Application of the HDAC inhibitors *in vitro* and *in vivo* enhances neuronal differentiation of progenitor cells, accompanied by an increased expression of different neuronal transcription factors (Hsieh et al., 2004; Siebzehnrubl et al.,2007; Yu et al., 2009). HDACs have been found to modulate not only cell differentiation, but also neuron specialization. Inhibition of HDACs activity negatively regulated the expression of genes responsible for GABA synthesis and the development of GABAergic inhibitory neurons in cortical neuron cultures (Fukuchi et al., 2009). On the contrary, inhibition of HDACs in organotypic raphe slice cultures stimulated the expression of genes responsible for serotonin synthesis and through the AMPAR-CaMKII signaling cascade enhanced its release (Asaoka et al., 2015).

Epigenetic rearrangements create a favorable environment for the intrinsic plasticity of brain cells (Ehrensberger and Svejstrup, 2012), where they are able to transform into cells of a different lineage (transdifferentiation or direct reprogramming) or to return to a pluripotent state (reprogramming), from where they can differentiate into virtually any cell type. Transdifferentiation has been observed in natural conditions including both pathological and normal developmental states. For instance, transdifferentiation occurs as a regenerative response to injuries: following brain injury, the mature striatal astrocytes are capable to transdifferentiate into functional cholinergic or GABAergic neurons (Duan et al., 2015); similarly, after the spinal cord injury the oligodendrocyte precursor cells, also known as NG2 glia, can generate excitatory and inhibitory propriospinal neurons (Tai et al., 2021). In both cases, the newly generated neurons successfully integrate into existing neural networks. The oligodendrocyte progenitors and reactive astrocytes represent the most often targets for cellular reprogramming due to their high lineage plasticity among other brain cells (Niu et al., 2013; Boshans et al., 2019; Pavlou et al., 2019; Griffiths et al., 2020). Conversion of glia into the subtype-specific neurons can be induced both *in vitro* and *in vivo* under certain experimental conditions aimed at changing the expression of proneural genes by transcription factors (Torper et al., 2013; Lentini et al., 2021), microRNAs (Weinberg et al., 2017; Mo et al., 2018), and small molecules cocktails (Qin et al., 2017; Ma et al., 2021; Pavlova et al., 2022). Further unraveling of the molecular pathways underlying brain cell identity may offer new strategies for regenerative medicine and novel therapeutic approaches to repair or replace malfunctioning cells in neurological diseases such as Alzheimer’s disease, epilepsy, etc (Yavarpour-Bali et al., 2020; Lentini et al., 2021; Giehrl-Schwab et al., 2022).

In the current study, we assessed how chromatin rearrangements, induced by broad-spectrum HDAC inhibitors, affect cell proliferation and differentiation in rat primary cultures using transcriptomics, qPCR, click-chemistry and immunocytochemistry. We performed a complex characterization of the epigenetically regulated neuronal and glial transcriptional programs from the viewpoint of cell differentiation and evaluated the expression of several cell-specific markers. HDAC inhibitors were found to attenuate cell proliferation while promoting the expression of various master transcription factors involved in cell differentiation and specialization. The number of DAPI+ cells remained unchanged after treatment, but some cells apparently underwent qualitative changes in cellular phenotype. Particularly, we revealed a significant upregulation of genes, normally expressed in neuromodulatory neurons (serotonergic, dopaminergic, cholinergic), and downregulation of genes, expressed in glial cells and inhibitory neurons. Transcriptional changes were accompanied by elevation of histone serotonylation levels both in neurons and in populations of glial cells. Based on these observations, we hypothesized that the detected posttranslational histone modifications not only regulate cell differentiation, but could be potentially involved in the regulation of cell identity and cellular reprogramming.

## Results

### HDAC inhibitors shift the gene expression profiles in primary neuron cultures from proliferation to differentiation and affect the expression of master transcription factors

To evaluate how epigenetic rearrangements affect gene expression profiles in brain cells, we performed bulk RNA sequencing of samples from rat primary neuron cultures treated with broad-spectrum histone deacetylase (HDAC) inhibitor trichostatin A (TSA, 100 nM) as previously described (Borodinova et al., 2019). In addition, to exclude possible non-specific actions of TSA on gene expression profile, we tested another broad-spectrum HDAC inhibitor sodium butyrate (NaB, 5 µМ) applied for 24 hours. Transcriptome analysis revealed significantly overlapping datasets of 6431 and 6347 differentially expressed genes (DEGs) in cortical neuron cultures treated with TSA or NaB, respectively (Fig.1A) (Supplemental_Table_S1). Heatmap clustering of 4930 overlapping DEGs revealed similar transcriptional profile for TSA-treated and NaB-treated cultures that differed from controls (Fig.1B). A significant number of the overlapping DEGs in the tested groups suggests that these HDAC inhibitors with different chemical structures share common regulatory pathways directly related to the regulation of transcriptional programs in primary neuron cultures.

**Fig. 1.**
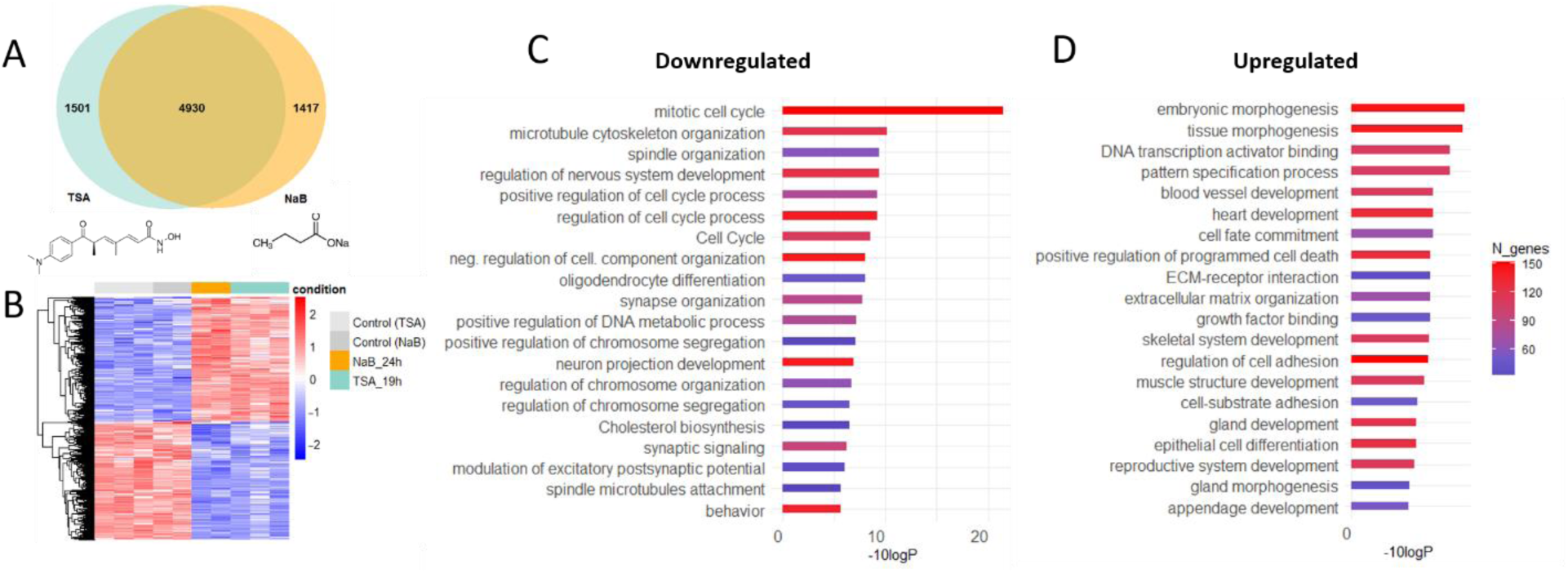
HDAC inhibitors TSA and NaB induce transcriptional changes in primary neuron cultures. A – Venn diagram illustrates the number of unique and overlapping differentially expressed genes (DEGs) in TSA-treated and NaB-treated neuron cultures. B - Cluster analysis of overlapping DEGs. Heatmap of 4930 overlapping DEGs shows differential expression in NaB-treated (orange) and TSA-treated neuron cultures (turquoise) compared to control cultures (gray). The color scale indicates the expression levels (blue, low expression; red, high expression). C,D - Gene Ontology (GO) analysis of the main enriched genes after HDAC inhibitors treatment. Bar charts show the top 20 most enriched GO terms for downregulated (C) and upregulated DEGs (D) based on Metascape. Gene ontologies are ranked by their significance.

Overlapping DEGs were represented by 2401 downregulated and 2529 upregulated genes (Supplemental_Table_S1). Gene Ontology (GO) analysis of DEGs datasets using Metascape software (Zhou et al., 2019) revealed that a significant part of downregulated genes is involved in biological processes related to cell organization and proliferation (Fig.1C), while upregulated genes were engaged in cell differentiation and specialization, tissue and embryonic morphogenesis (Fig.1D). The clusters of downregulated genes were associated with cell division and DNA replication, including different cyclins and cyclin-dependent kinases (Fig.2A). To verify the influence of HDAC inhibitors on cell proliferation, we pretreated cell cultures for 1 hour with 10 μM EdU that incorporates into newly synthesized DNA of proliferating cells. Subsequent analysis revealed a significant reduction in the number of EdU-positive cells in primary neuron cultures treated with TSA, reflecting a decrease in the proliferation rate (Fig.2B).

**Fig. 2.**
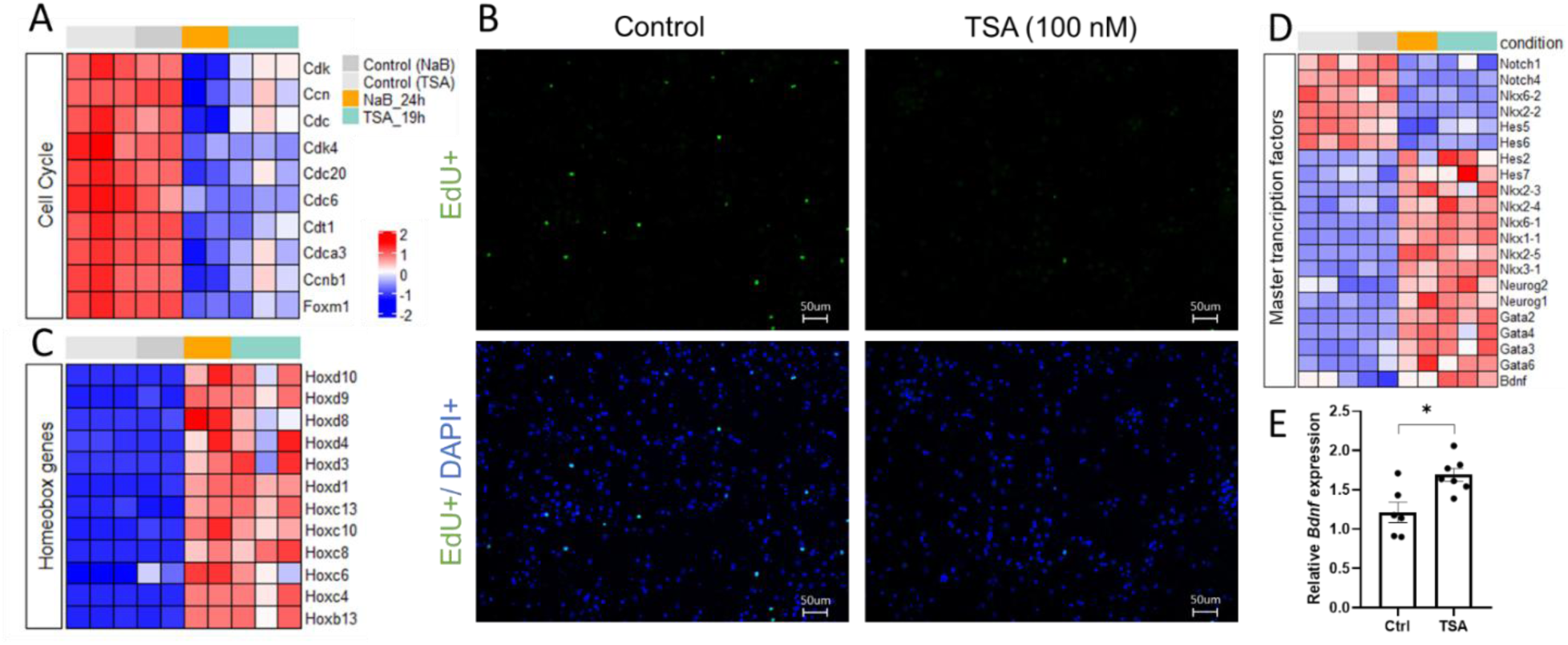
HDAC inhibitors impair cell proliferation and stimulate cell differentiation in rat primary neuron cultures. A – Heatmap illustrates representative downregulated genes associated with cell cycle. B – Representative micrographs of EdU-positive proliferating cells (green) in control and TSA-treated primary neuron cultures, revealed by click-chemistry. Cell nuclei are labeled with DAPI (blue). Scale bar was 50 µM. Magnification 20x. C-D – Heatmaps illustrate representative upregulated homeobox genes (C) and master transcription factors (D), which control embryonic morphogenesis and brain cells differentiation. The color scale indicates the expression levels (blue, low expression; red, high expression). E –Quantification of the relative mRNA expression of differentiation factor BDNF using qPCR. * p < 0.05.

In the DEGs datasets, we found distinct classes of both downregulated and upregulated master transcription factors that work together to control embryonic morphogenesis, brain cell differentiation and specialization (Fig.2C,D). Among the downregulated genes, we identified several transcription factors that control glial cell differentiation (Nkx6.2, Nkx2.2, Notch1, etc) (Pavlou et al., 2019; Sock and Wegner, 2021) (Fig.2D). In a dataset of upregulated genes, we identified proneural transcription factors from different families of homeobox proteins (Hox, Nkx), basic helix–loop–helix proteins (Neurog1, Neurog2), zinc finger DNA binding proteins (Gata), and neurotrophins (Bdnf, NT4) (Fig.2C,D), known evolutionarily conserved master regulators of embryonic morphogenesis that form the neurogenic axis of cellular differentiation and promote neuron specialization (Rumajogee et al., 2002; Craven et al., 2004; Dixit et al., 2014; Seifert et al., 2015; Haugas et al., 2016; Shimojo et al., 2024). Using qPCR we confirmed that mRNA expression of brain-derived neurotrophic factor BDNF, the inducer of brain development and plasticity, was significantly upregulated in response to application of HDAC inhibitors (Fig.2E).

### HDAC inhibitors regulate the expression of specific markers of various brain cells

Given that treatment with HDAC inhibitors suppresses cell proliferation and regulates the expression of specific master transcription factors involved in cell differentiation, we sought to characterize clusters of epigenetically controlled genes associated with differentiation and specialization of individual brain cells. In a diagram on Fig.3 we have combined publicly available data on the generally accepted unique markers of differentiating and mature brain cells (https://www.cellsignal.com/pathways/neuronal-and-glial-cell-markers; https://www.abcam.com/neuroscience/neural-markers-guide) with the results of our transcriptome analysis of downregulated (blue) and upregulated (red) differentially expressed genes (DEGs) from overlapping datasets (Fig.3).

**Fig. 3.**
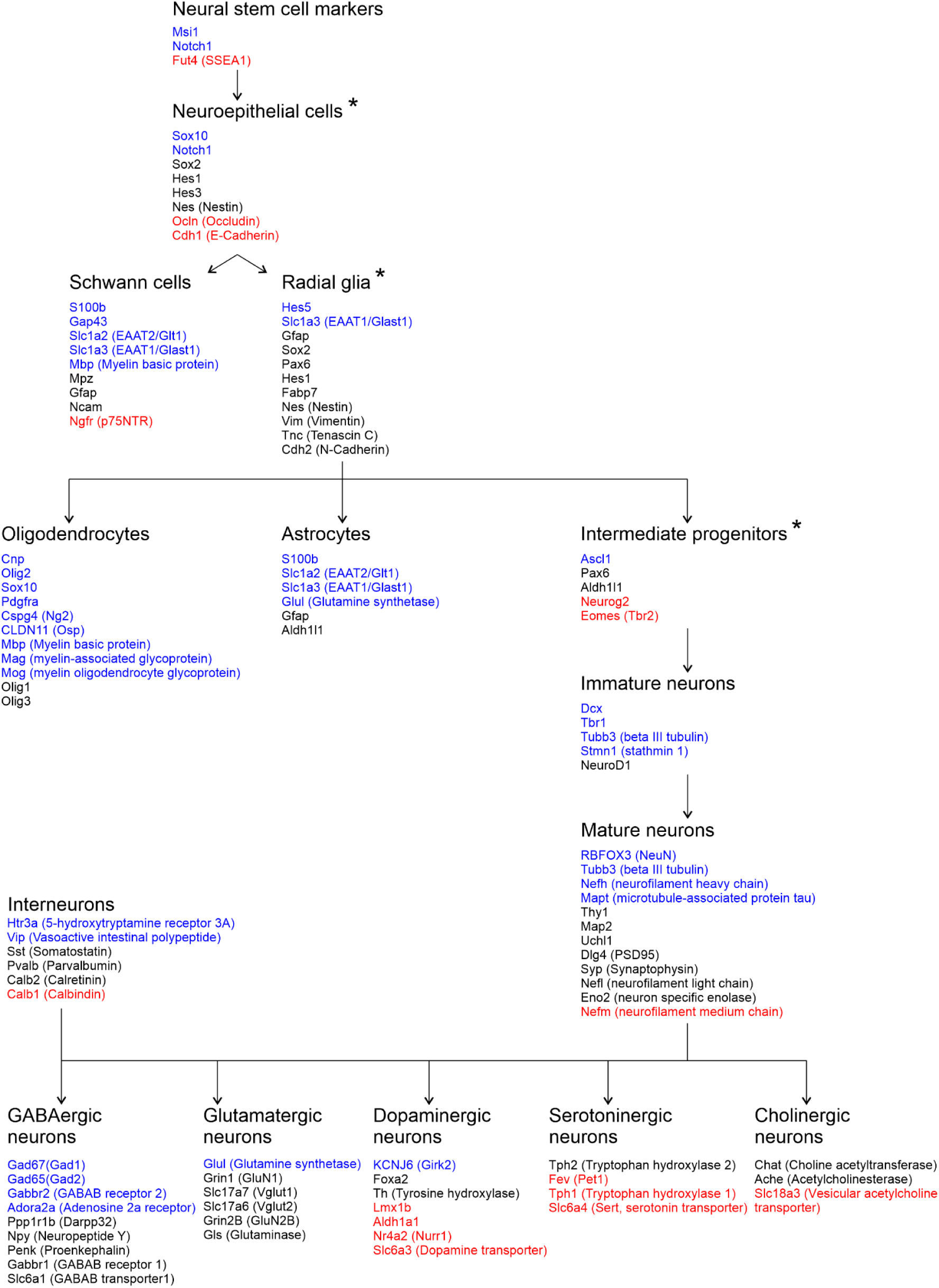
Scheme shows specific markers of diverse brain cells at different stages of their differentiation and specialization. Genes, whose expression increases (red) or decreases (blue) after treatment of rat primary neuron cultures with histone deacetylase inhibitors, are marked on the diagram. * labels the proliferating cells.

We found that transcription of multiple glial and neuronal markers was affected in rat primary neuron cultures in response to application of the HDAC inhibitors NaB or TSA. Particularly, the transcriptome analysis revealed a profound loss of unique glial markers typical for astrocytes (Fig.4A), as well as for oligodendrocytes and oligodendrocyte precursor cells (Fig.4B). The RNA sequencing data were confirmed by quantitative PCR using pairs of primers for several astrocyte (*Aqp4*, *Slc1a2*, Fig.4C) and oligodendrocyte markers (*Sox10*, *Opalin*, *Olig2*, Fig.4D). These results are consistent with previously published data, showing the requirement of HDACs for oligodendrocytes differentiation, while less is known about their involvement in astrocytes differentiation (Humphrey et al., 2008; Ye et al., 2009; Conway et al., 2012; Castelo-Branco et al., 2014; Zhang et al., 2016).

**Fig. 4.**
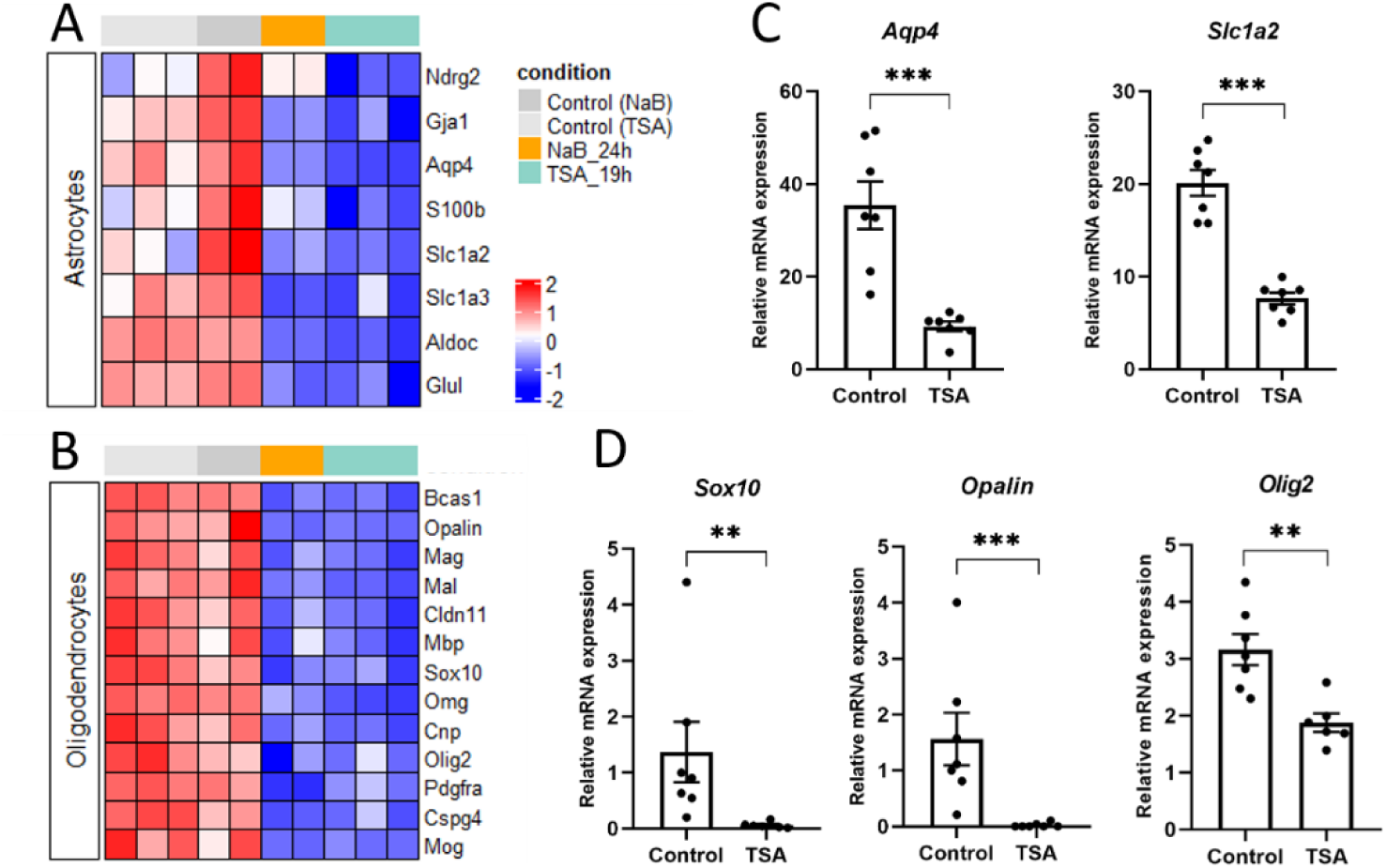
HDAC inhibitors alter mRNA expression of specific glial cell markers. A,B - Heatmaps illustrate representative downregulated genes encoded specific astrocyte markers (A) and oligodendrocyte markers related to myelin synthesis process (B). The color scale indicates the expression levels. C,D – qPCR verification of RNAseq data using specific primer pairs for astrocyte (C) and oligodendrocyte markers (D). * p < 0.05.

mRNA expression of the main neuron-specific markers such as NeuN (Rbfox3), beta III tubulin (Tubb3) and microtubule-associated protein tau (Mapt) was affected in our experimental conditions based on transcriptome analysis (Fig.3) and quantitative PCR (Fig.5A). Interestingly, with a general decline in the expression of neuronal markers, transcription of specific markers of different neuronal subpopulations was regulated in opposite ways (Fig.5B). Similarly to previous data (Fukuchi et al., 2009), we found that the expression of interneuron markers, associated with GABA synthesis, transport and signaling (*Gad1*, *Gad2*, *Gabbr2, Slc6a11*), was significantly downregulated after application of HDAC inhibitors (Fig.5 B,C). Since the expression of *Htr3a* and *Vip* genes was decreased upon induction of epigenetic rearrangements (Fig.5C), we hypothesized that the most profound transcriptional changes affected the Htr3a+/VIP+ interneuron population, which constitutes approximately one-third of cortical interneurons (Rudy et al., 2011).

**Fig. 5.**
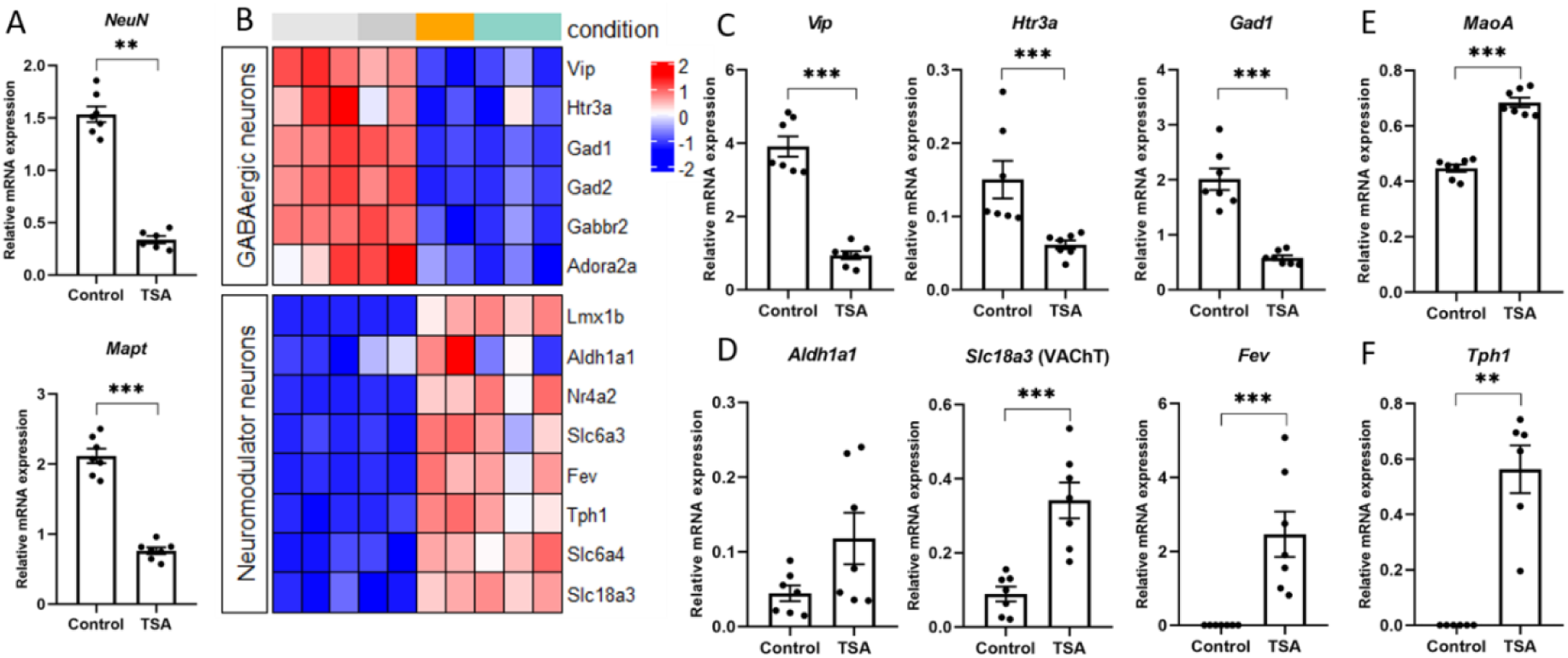
HDAC inhibitors influence mRNA expression of neuronal markers. A – qPCR verification of RNAseq data using specific primer pairs for neuronal markers. B – Heatmap illustrates representative downregulated and upregulated genes specific for GABAergic interneurons (top panel) and neuromodulatory neurons (bottom panel). The color scale indicates the expression levels (blue, low expression; red, high expression). C – qPCR verification of RNAseq data using specific primer pairs for *Htr3a*, *Vip*, and *Gad1* genes specific for GABAergic interneurons. D – qPCR verification of RNAseq data using specific primer pairs for genes, specific for distinct neuromodulatory neurons, including *Aldh1a1* (dopaminergic), *Slc18a3* (cholinergic) and *Fev* (serotonergic). E,F – qPCR analysis of the expression of *Tph1* and *MaoA* genes encoding crucial enzymes for serotonin metabolism. * p < 0.05.

Interestingly, HDAC inhibitors positively regulate the expression of genes associated with secretory phenotypes of neuromodulatory neurons (Bence et al., 2012; Asaoka et al., 2015). Indeed, in our experiments on primary neuron cultures the expression of genes encoding transcription factors (*Fev*, *Lmx1b*, *Nr4a2*), enzymes (*Tph1*, *Aldh1a1*, *MaoA*) and transporters (*Slc6a4*, *Slc6a3*, *Slc18a3*), specific for distinct neuromodulatory neurons, was significantly upregulated in response to epigenetic rearrangements induced by the HDAC inhibitors (Fig.3, Fig.5B). Using qPCR, we confirmed that application of HDAC inhibitors stimulated the expression of several serotonergic (*Fev* aka Pet1, *Tph1*) and cholinergic genes (*Slc18a3* aka VAChT), but not the dopaminergic genes (*Aldh1a1*) (Fig.5D,F). Presumably, this effect is directly related to regulation of gene transcription. However, the known indirect neuroprotective effects of HDAC inhibitors on the survival of neuromodulatory neurons and the maintenance of their phenotype should be noted (Chen et al., 2006).

Thus, our results in primary neuron cultures, although consistent with previously published data, additionally demonstrate multiple parallel trajectories of differentiation/de-differentiation processes simultaneously driven by epigenetic rearrangements.

### HDAC inhibitor trichostatin A elevates the expression of genes associated with serotonergic secretory phenotype and stimulates histone serotonylation

During detailed analysis of the upregulated DEGs dataset we revealed a subset of genes encoding important components of the serotonergic cells’ functioning including the 5-HT transporters (*Slc6a4* aka Sert; *Slc18a1* aka VMAT1), and enzymes for the synthesis (*Tph1*) and degradation of 5-HT (*Maoa*). To verify the RNA sequencing data, we performed qPCR with specific primer pairs for the *Tph1* (Fig.5F) and *Maoa* (Fig.5E) genes, encoding key enzymes responsible for the 5-HT turnover (Fig.6A), and confirmed that application of TSA significantly increases the expression of target genes in both neurons and glia (Supplemental_Fig_S1A). These results are consistent with previously published data on serotonergic neurons and non-neuronal cells that describe HDACs as negative regulators of serotonergic phenotype and demonstrate positive influence of their inhibitors on the expression of different genes involved in serotonin turnover (Asaoka et al., 2015; Phi van et al., 2015; Zhang et al., 2020). In these studies, activation of serotonin-associated genes was accompanied by the accumulation, release and uptake of 5-HT in cultured cells. Potentially, similar effects could be observed in our experimental conditions.

**Fig. 6.**
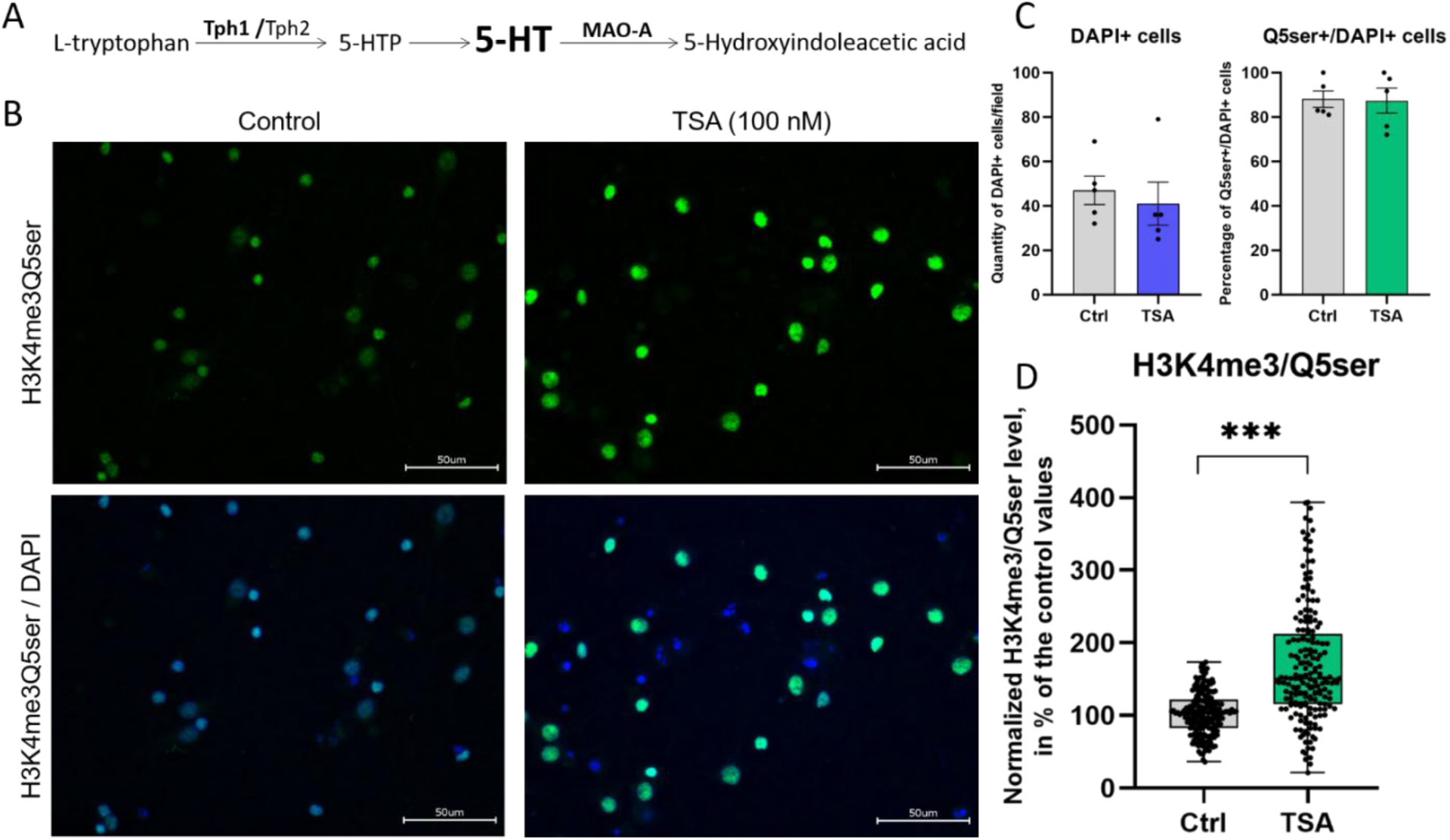
HDAC inhibitor trichostatin A (TSA) enhances histone serotonylation in rat primary cortical cultures. A – Scheme of serotonin (5-HT) synthesis and metabolism with indication of key enzymes; B – Fluorescent microscopy of cell cultures stained with antibodies against H3K4me3Q5ser that targets histone H3 trimethylated on lysine 4 and serotonylated on glutamine 5 (green), and DAPI (blue). Scale bar was 50 µM. Magnification 60x. C – Quantification of DAPI+ cells per field and percentage of H3K4me3Q5ser+ cells relative to total number of DAPI+ cell in control and TSA-treated cultures. Results are presented as Mean ± s.e.m. D – The box plot shows the results of densitometric analysis of histone serotonylation levels in control and TSA-treated cultures. Data were normalized and calculated as a % of control values in each biological replica, each point shows the value for the individual cell, n=5. Results are presented as Median ± SD. * p < 0.05.

Given that the 5-HT synthesis enzyme Tph1 can be expressed in non-neuronal cells, we examined its distribution both in neurons and in glia in primary cultures. Our preliminary results confirmed that Tph1 is widely expressed in differentiating cultured brain cells (Supplemental_Fig_S1C), providing them with the ability to synthesize serotonin.

Next, we questioned what are the functions of serotonin in differentiating cell cultures. It is well known that 5-HT acts as a classical neurotransmitter – modulator of synaptic processes, but also it acts as a factor involved in cell differentiation (Lavdas et al., 1997). Recent studies suggest that effects may occur through histone serotonylation – the posttranslational histone modification, when serotonin covalently binds to glutamine residues on histone proteins by the endogenous tissue transglutaminase 2 (TGM2), and alters gene transcription through epigenetic rearrangements (Farrelly et al., 2019). According to our transcriptomic data, *Tgm2* gene expression was predicted to be increased by the HDAC inhibitors. We hypothesized that, if true, it may contribute to the serotonylation of histones in primary cultures of cortical neurons treated with HDAC inhibitors. To test this hypothesis, cortical neuron cultures were stained with antibodies that recognize the previously described dual permissive mark H3K4me3Q5ser on histones that combines serotonylated glutamine at position 5 (Q5ser) on histone H3 and its neighboring trimethylated lysine at position 4 (H3K4me3) (Farrelly et al., 2019). We observed widespread expression of H3K4me3Q5ser mark in both control and TSA-treated primary neuron cultures, more than in 80% of all cells (Fig.6B,C). Taking into account that the percentage of DAPI+ cells expressing the histone serotonylation mark H3K4me3Q5ser was not changed (Fig.6C), a significant elevation of histone serotonylation in TSA-treated cells was confirmed by densitometric analysis of images (Fig.6D). We noticed that the total number of DAPI+ cells was unchanged (Fig.6C), which implies that the enhanced histone serotonylation was not due to changes in cellular quantity and composition. We conclude that HDAC inhibitors stimulate the expression of serotonergic genes in primary cultures of cortical neurons, which ensure the production of 5-HT and serotonylation of histones in the cells, potentially involved in changes in cell differentiation.

### Histone serotonylation marks are widely distributed across different populations of both neurons and glia

Widespread expression of the H3K4me3Q5ser mark raises the question of which cell populations undergo changes in histone serotonylation following HDACs inhibition. Immunocytochemical analysis revealed substantial overlap of the H3K4me3Q5ser mark with the main neuronal (Dcx, NeuN) and glial markers (GFAP, Olig2) (Fig.7A). We quantified a percentage of H3K4me3Q5ser+ cells that express cell-specific markers in control and TSA-treated groups and found no differences (Fig.7B-D, left panel). However, TSA treatment caused a significant increase in histone serotonylation levels in all double-labeled cells tested, including immature Dcx^+^ and mature NeuN^+^ neurons, as well as Olig2-positive oligodendrocytes (Fig.7B-D, middle panel). These changes were accompanied by a significant decrease in the mRNA (Fig.4D, Fig.5A) and protein expression of transcription factors NeuN and Olig2 (Fig.7C,D right panel), responsible for cell specialization.

**Fig. 7.**
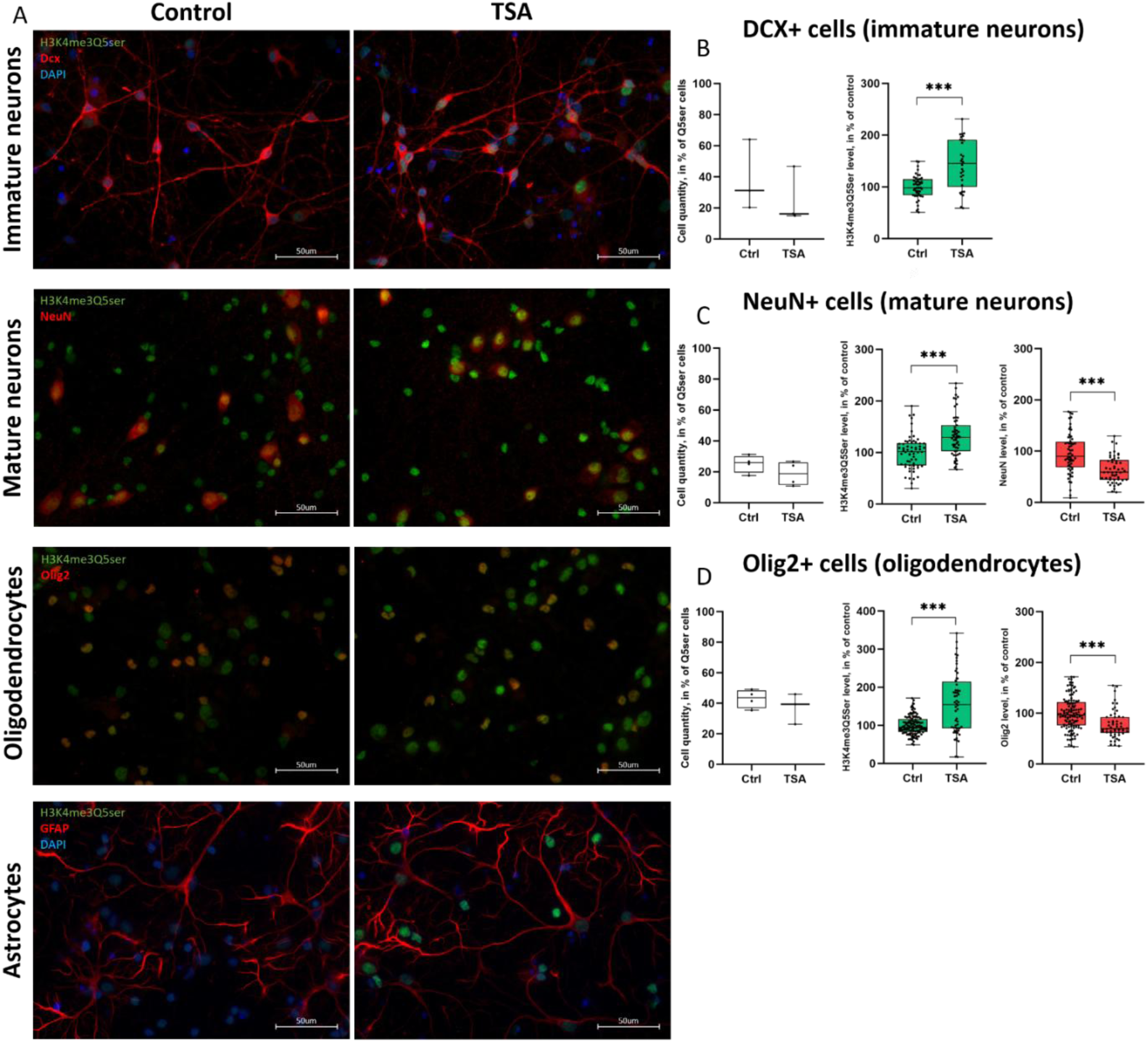
Various populations of brain cells were affected by HDAC inhibitor trichostatin A (TSA). A - Fluorescent microscopy shows co-localization of H3K4me3Q5ser marks (green) with one of the cell-specific markers (red), and DAPI (blue). Scale bar was 50 µM. Magnification 60x. B-D – Figures show the ratio of double-labeled cells relative to the total number of H3K4me3Q5ser+ cells (left panel); Densitometric analysis of histone serotonylation levels (middle panel) and the levels of cell-specific transcription factors in control and TSA-treated cultures. Data were normalized and calculated as a % of control values, each point shows the value for the individual cell. Results are presented as Median ± SD. * p < 0.05.

Since the elevated levels of histone serotonylation were detected in both neurons and glial cells, we wondered whether this was a cause or a consequence of widespread transcriptional changes induced by HDACs inhibitors. The time-course experiments showed early downregulation of gene expression (*Olig2, Sox10*) (Supplemental_Fig_S2A) along with increased histone serotonylation in cell cultures (Supplemental_Fig_S2B). The results obtained indicate that transcriptional changes were accompanied by increased histone serotonylation, which appeared after 4 hours of incubation with TSA and persisted for at least 19 hours. Given that histone serotonylation was increasing in distinct brain cells along with changing expression of cell-specific transcription factors, we speculate that chromatin remodeling induced by histone serotonylation contributes to the changes in a transcriptional program associated with cell fate commitment and cell specialization, but it requires further investigation.

## Discussion

Studying the principles of regulation of cell fate commitment is important for understanding the mechanisms of normal physiological processes as well as neurodegenerative diseases, when certain cells lose or change their identity and become abnormal. Epigenetic modifiers of the chromatin landscape play a crucial role in the regulation of transcriptional programs essential for establishment and maintenance of cellular phenotype. Changes in the activity of various epigenetic modifiers influence chromatin compaction and DNA accessibility to transcription factors, creating a favorable environment for the intrinsic plasticity of brain cells (Ehrensberger and Svejstrup, 2012). In the current study, we aimed to determine how chromatin rearrangements, induced by broad-spectrum HDAC inhibitors, affect cellular differentiation trajectories. Unlike previous studies on specialized cell lines or multipotent stem/progenitor cells, we used differentiating rat primary neuron cultures to describe epigenetic regulation of transcriptional programs in distinct brain cells using RNA sequencing. To eliminate possible side effects and to increase the reproducibility and reliability of our results, we tested two different HDAC inhibitors with different chemical structures in parallel. Trichostatin A (TSA) and sodium butyrate (NaB) produced very similar transcriptional changes (Fig.1A,B, Supplemental_Table_S1) that include downregulation of genes involved in cell organization and proliferation (Fig.1C, Fig.2A), glial fate commitment and glial differentiation (Fig.2D, Fig.4). The reduction in the proliferative potential of primary cultures was confirmed by click-chemistry with EdU, which incorporates into newly synthesized DNA of proliferating cells (Fig.2B). These results are consistent with previous findings demonstrating that HDACs are required for cell proliferation and neurogenesis (Jiang and Hsieh, 2014; Tang et al., 2019) while their inhibition with small molecules (TSA, NaB, valproic acid) attenuate significantly cell proliferation, stimulate the expression of proneural transcription factors and promote neuronal differentiation both *in vitro* and *in vivo* (Hsieh et al., 2004; Yu et al., 2009).

In the overlapping DEGs datasets we identified distinct classes of master transcription factors that work together to control cell differentiation and specialization (Supplemental_Table_S1). Particularly, the expression of transcription factors promoting glial cell differentiation (Olig2, Sox10; Nkx6.2, Nkx2.2, Notch1, etc) (Pavlou et al., 2019; Sock and Wegner, 2021) was significantly downregulated as confirmed by transcriptome analysis (Fig.2D), qPCR (Fig.4) and ICC (Fig.7D). This was accompanied by a decrease in the expression of different glial markers, with the most pronounced suppression of oligodendrocyte genes (Fig.4B, Supplemental_Table_S1). HDAC activity was shown to be essential for glial differentiation and maturation, they interact with regulatory regions on chromatin and control specific transcriptional programs (Castelo-Branco et al., 2014). We know from the literature that distinct glial cells (oligodendrocytes, astrocytes) lose open chromatin marks during differentiation as compared to multipotent cells and neurons (Hsieh et al., 2004; Douvaras et al., 2016). In fact, HDACs associated with repressor complexes containing Sin3A or NcoR co-repressors (Borodinova, Balaban, 2020) are recruited to regulatory regions on chromatin to control glial cell identity through the repression of proneural gene activity in non-neuronal cells (Huang et al., 1999; Castelo-Branco et al., 2014). Additionally, HDACs can promote the expression of pro-glial factors (Castelo-Branco et al., 2014), likely through the previously described interaction with histone acetyltransferases (Zhang et al., 2016).

Numerous studies have demonstrated that chromatin remodeling induced by HDAC inhibitors prevents neural progenitor cells (NPCs) differentiation into glial cells and favors differentiation into neurons (Hsieh et al., 2004; Balasubramaniyan et al., 2006; Siebzehnrubl et al., 2007; Yu et al., 2009). In our transcriptomic data we observed an increased expression of proneural transcription factors from distinct families of homeobox proteins (Hox, Nkx), basic helix–loop–helix proteins (Neurog1, Neurog2), zinc finger DNA binding proteins (Gata), and neurotrophic factors (Bdnf, NT4) (Fig.2D, Fig.3), which form the neurogenic axis of cell differentiation (Shimojo et al., 2024), determine neural lineage commitment and promote neuronal differentiation and specialization (Rumajogee et al., 2002; Craven et al., 2004; Dixit et al., 2014; Haugas et al., 2016). Surprisingly, several markers of fully mature neurons (Mapt, NeuN) were significantly downregulated as confirmed by qPCR and ICC staining (Fig.3, Fig.5A,B, Fig.7C). Interestingly, some studies describe their heterogeneous expression in different subpopulations of neurons, as well as in glial cells (Darlington et al., 2008; Cannon and Greenamyre, 2009; Seiberlich et al., 2015). Therefore, we suggest that downregulation of “neuronal” markers, observed in our experiments, is in fact a reflection of significantly altered transcriptional programs underlying different cellular identities, which may lead to changes in cell composition, possibly through cellular reprogramming.

Heterogeneous by nature, distinct neuron subpopulations have a certain set of properties that are acquired through a certain transcriptional program, controlled by specific transcription factors. These transcriptional programs can be epigenetically regulated in different ways during neuron specialization (Fukuchi et al., 2009; Asaoka et al., 2015). In our transcriptomic data we found opposite regulation of genes specifically expressed in subpopulations of inhibitory neurons and neuromodulatory neurons. Particularly, we observed downregulation of multiple interneuron genes involved in GABA synthesis, transport and signaling following HDAC inhibitor treatment (Fig.5B,C), as previously shown (Fukuchi et al., 2009). Recent data suggest that cortical interneurons are represented by three populations of cells selectively expressing either parvalbumin (PV), somatostatin (SST), or ionotropic serotonin receptor 5HT3a (Htr3a), the latter, in turn, is divided into VIP-positive and VIP-negative subpopulations (Rudy et al., 2011). We confirmed the downregulation of *Htr3a* and *Vip* genes encoding interneuron markers upon induction of epigenetic rearrangements (Fig.5C). Given that, we assume a subset of cortical Htr3a^+^/VIP^+^ and/or Htr3a^+^/VIP^-^ interneurons were most likely affected, but it requires further investigation.

Interestingly, downregulation of the GABAergic genes in primary neuron cultures was counterbalanced by the upregulation of multiple genes associated with neuromodulatory neuron phenotypes (Fig.3). Among the upregulated DEGs, we identified master transcription factors (Gata2, Gata3, Fev aka Pet1, Lmx1b, Nr4a2 aka Nurr1) that are responsible for establishing and maintaining the phenotypes of serotonergic and dopaminergic neurons (Craven et al., 2004; Krueger and Deneris, 2008; Hoekstra et al., 2013; Haugas et al., 2016). This was accompanied by elevated expression of the corresponding cell-specific markers, particularly, the important components of serotonin turnover (Fig.5B, D-F). We confirmed that the serotonin synthesis enzyme Tph1 is widely expressed in cortical neurons and astrocytes of primary neuron-glia cultures (Supplemental_Fig_S1C), and its transcription is elevated by the HDAC inhibitors (Fig.5F, Supplemental_Fig_S1A), which may increase the ability of cells to synthesize serotonin. These results are in agreement with previous studies describing 1) the negative influence of HDACs on the serotonergic phenotype and 2) the positive influence of HDAC inhibitors on serotonergic gene expression accompanied by increased serotonin synthesis and release (Asaoka et al., 2015; Phi van et al., 2015; Zhang et al., 2020).

It is well established that serotonin is a multifaceted regulator of many biological processes in the brain, acting in synapses as a neurotransmitter and in the nucleus as a chromatin remodeling factor. Serotonin serves as a donor of monoamine groups for posttranslational modification of various proteins called serotonylation, catalized by tissue transglutaminase 2 (TGM2) (Rossin et al., 2023). Recent studies have demonstrated that TGM2 mediates serotonylation of glutamine residues on histone proteins, leading to stabilization of adjacent marks H3K4me3 and epigenetic regulation of gene expression programs essential for brain development and functioning (Farrelly et al., 2019; Zhao et al., 2021; Sardar et al., 2023; Al-Kachak et al., 2024). According to our transcriptomic data, the expression of *Tgm2* gene was upregulated after application of HDAC inhibitors. We hypothesized that this might stimulate histone serotonylation under our experimental conditions. Indeed, immunocytochemical staining revealed increased levels of histone serotonylation in TSA-treated primary cortical cultures (Fig.6B,D). Elevation of histone serotonylation was widely distributed across different populations of both neurons and glial cells as confirmed by co-localization with various cellular markers (Fig.7). We found that temporal dynamics of gene expression in TSA-treated cell cultures (Supplemental_Fig_S2A) was accompanied by increased histone serotonylation, which appeared after 4 hours of incubation with TSA and persisted for at least 19 hours (Supplemental_Fig_S2B; Fig.6 B).

We hypothesized that the observed switching of transcriptional programs associated with phenotypically distinct brain cells indicates the onset of cell fate changes, possibly through the initiation of reprogramming processes. Previous studies demonstrated that induction of neuronal fate conversion both *in vitro* and *in vivo* can be achieved by (epi)genetic manipulations with a single transcription factor or a panel of transcription factors in glial cells (astrocytes, oligodendrocyte precursor cells) (Lyssiotis et al., 2007; Niu et al., 2013; Torper et al., 2013; Pereira et al., 2019; Lentini et al., 2021). Also, some neuronal subpopulations were found to change their secretory phenotypes following genetic manipulation with transcription factors (Raina et al., 2020). Importantly, the reprogramming efficiency and complete maturation of differentiated neurons depend on a permissive microenvironment created by chromatin remodeling and neurotrophic factors (Balasubramaniyan et al., 2006; Niu et al., 2013; Rivetti di Val Cervo et al., 2017).

These data support our hypothesis that the observed complex transcriptional changes, involving a number of transcription factors and neurotrophins, most likely contributed to onset of cellular reprogramming under our experimental conditions. We hypothesized that HDACs inhibitors induce chromatin remodeling by creating open chromatin marks that disrupt existing steric barriers to Tgm2 activity (Lukasak et al., 2022), and facilitate histone serotonylation in regulatory regions of target genes. This results in serotonylation of glutamine residues at position 5 (Q5ser) on histone H3 and stabilization of adjacent trimethylated lysine at position 4 (H3K4me3) by creating a dual permissive mark H3K4me3Q5ser that facilitates gene transcription through interaction with the transcription initiation factor TFIID (Farrelly et al., 2019). The early appearance of enhanced histone serotonylation marks and the persistence of these changes over many hours in distinct brain cells may indicate that chromatin remodeling induced by histone serotonylation contributes to the maintaining of a new transcriptional programs associated with cell fate commitment and cell specialization, but this requires further investigation.

## Materials and methods

### Animals

The experiments were carried out in newborn (P0–P1) Wistar rats (Pushchino breeding facility, Russia). All experimental procedures were conducted in accordance with the European Communities Council Directive of 24 November 1986 (86/609/EEC) on the protection of animals used for scientific purposes. The study protocol was approved by the Ethics Committee of the Institute of Higher Nervous Activity and Neurophysiology of RAS.

### Rat primary cortical neuron cultures

Cell cultures were prepared as previously described (Borodinova et al., 2019). For qPCR experiments, approximately 0.25-0.3 million of isolated cells were plated into individual wells coated with poly-D-lysine hydrobromide (Sigma). For ICC experiments, the same number of cells was placed in individual wells on 12 mm glass coverslips coated with poly-D-lysine hydrobromide. Primary cortical neuron cultures were grown for two weeks in CO2 incubator (5% CO2, 37 °C), culture medium was partially refreshed every 2-3 days. HDAC inhibitors were applied on the 14th day *in vitro* (DIV) without washout.

### Drugs

Two broad-spectrum histone deacetylase (HDAC) inhibitors with different chemical structures were used in the study. The HDAC inhibitor trichostatin A (TSA, 100 nM, Sigma) was applied for 19 hours as previously described (Borodinova et al., 2019). The HDAC inhibitor sodium butyrate (NaB, 5 µМ, Sigma-Aldrich) was applied for a longer time of 24 hours.

### Cell Proliferation Assay

Proliferating cells were identified by labeling replicated DNA with ethynyl deoxyuridine (EdU, Lumiprobe). Following its incorporation into DNA, EdU was subsequently detected with a fluorescent azide (Alexa Fluor™ 488 Azide, Molecular Probes) via “click” chemistry reaction.

EdU (10 μM, Lumiprobe) was added to half of the culture medium for 1 hour before drug administration. After incubation with EdU, the medium was replaced with another half of the medium supplemented with the HDACs inhibitor TSA. After 19 h of incubation with TSA, cultures were fixed for 10 min at room temperature (RT) in 4% PFA and permeabilized for 15 min with 0.1% Triton X-100 in PBS. EdU-labeled cells were then stained for 30 min with a “click” reaction mixture (4 mM copper(II)-BTTAA complex, 100 mM ascorbic acid, 12 μM dye azide in 100 mM Tris buffer, pH 8.5).

### RNA extraction and sequencing

Total RNA from primary neuron cultures was isolated using ExtractRNA kit according to the manufacturer’s protocol (Evrogen, Moscow, Russia). The following library preparation and sequencing was performed at Genomed company (Russia). Briefly, mRNA was purified from a total RNA mixture using oligo(dT)-coated magnetic beads, followed by mRNA fragmentation and cDNA synthesis. The synthesized cDNA was then subjected to end repair, 3’-adenylation and adapter ligation. Following amplification of cDNA fragments, the PCR products were purified using Ampure XP Beads (AGENCOURT) and dissolved in EB solution. Library was validated on the Agilent Technologies 2100 bioanalyzer. The double-stranded PCR products were then heat denaturated and circularized using splint oligo sequence to generate a final library of single-stranded circular DNA molecules. The library was amplified using phi29 DNA polymerase to create a DNA nanoball containing more than 300 copies of a single DNA molecule. Paired-end sequensing of the library was performed on the DNBseq-G400 platform at Genomed company (Russia).

### RNA sequence alignment and analysis

High quality reads were mapped onto the reference Rat genome (Rnor6, ENSEMBL database) using STAR (Dobin et al., 2013) with default parameters and raw read counts obtained using FeatureCounts (Liao et al., 2014). For each dataset, counts were normalized by “median-of-ratios” method and subjected to differential expression analysis using DESeq2 R-package (Love et al., 2014) with chosen p.adjusted significance level < 0.05.

### Visualization and functional annotation of DEGs

Regularized logarithm expression values were used for constructing heatmaps. Values on the heatmaps were presented as z-scored normalized expression counts. Gene ontology enrichment of differentially expressed genes was performed using Metascape web-resource (Zhou et al., 2019). Gene expression visualizations were done using R-packages ggplot2, ComplexHeatmap, VennDiagramm.

### qPCR analysis

Verification of sequencing data was performed using quantitative real-time PCR (qPCR). Equal amount of total RNA from each sample (approximately 800 ng) was treated with DNaseI (Thermo Scientific, MA, USA) and then taken for the first-strand cDNA synthesis using MMLV RT kit (Evrogen, Moscow, Russia) and random decamer primers (Evrogen). To verify sequencing data we performed qPCR using SYBR-green mastermix reagent (Evrogen) and specific primer pairs (Supplemental_Table_S2). For each sample, the reaction was run in triplicates in 384-well plates on the C1000 Touch Thermal Cycler (Bio-Rad, CA, USA) according to the following cycling conditions: 95 °C for 5 min; 42 cycles of 95 °C for 30 s, 61 °C (60 °C for *Htr3a*) for 30 s and 72 °C for 30 s. Values were normalized to the rat *Hprt* housekeeping gene (Supplemental_Table_S2). Relative mRNA expression was calculated with standard ΔΔCt method.

qPCR data are presented as Mean ± s.e.m. Statistical significance of differences between the groups was calculated with Mann-Whitney U test. Significance was set at p < 0.05. All qPCR experiments were performed in at least four biological replicas.

### Immunocytochemistry (ICC)

Control and TSA-treated cultures of primary rat cortical neurons were fixed for 10 min with 4% paraformaldehyde, washed with PBS, and then permeabilized for 15 min with 0.1% Triton X100. Depending on the secondary antibodies used the cells were blocked in 5% normal goat serum (NGS) or in 5% donkey serum dissolved in PBS for 1h at room temperature. Then cells were incubated with primary antibodies (Supplemental_Table_S3) for 1h at room temperature and washed three times for 10 min with 0.05% Tween20 in PBS. Secondary antibodies were used under the same conditions. DAPI (1:1000) was applied for 10 min and then washed with PBS. Coverslips were mounted with self-made Mowiol mounting medium supplemented with 8% DABCO. When the dilution of antibodies ranged from 1:50 to 1:200, we performed immunostaining on the parafilm in a small volume of 30 mkl by inverting the coverslips with the cells facing down onto the antibody drop, covering with a lid to prevent air drying, and incubating for 1h at room temperature. The coverslips were then returned to the plate and processed as usual. Quantitative ICC experiments were performed in at least three biological replicates.

### Microscopy and analysis

Microscopy was carried out using equipment of the Research Resource Center of IHNA and NPh RAS for functional brain mapping. Stained cell cultures were visualized using a fluorescence microscope (HS All-in-one Fluorescence Microscope BZ-9000, Keyence, USA) with 10x/0.45 numerical aperture (NA), 20x/0.75 NA, or 60x/1.40 NA (oil-immersion) Plan Apo λ objectives (Nikon). Z-stack images were acquired using the BZ-II Viewer with identical exposure parameters applied separately to each channel for each pair of control and experimental coverslips. Subsequent image preparation and analysis were performed using ImageJ software (NIH). Densitometric analysis involved manually selecting stained cells using the ROI Manager tool and creating a set of ROIs for each fluorescence channel, followed by measuring the fluorescence intensity in a single image from z-stack and normalizing to the background fluorescence intensity averaged over five areas. Data were presented as Median ± SD in % of control values. Statistical significance of differences between the groups was calculated with Mann-Whitney U test. Significance was set at p < 0.05.

## Acknowledgments

We thank Dr. Alexander Lazutkin (IHNA&NPh RAS) for providing EdU for analysis of cell proliferation. We thank Alexander Moshchenko (FCBRN FMBA, Russia) for providing the primary anti-GFAP antibodies.

## Author Contributions

Conceptualization, A.A.B. and A.P.B., investigation, A.A.B., Yu.A.L., and A.P.B.; data curation, A.V.R. and G.V.P.; writing—original draft preparation, A.A.B. and Yu.A.L.; writing—review and editing, P.M.B., and G.V.P.; supervision, P.M.B; funding acquisition, A.A.B. All authors discussed the results and contributed to the final manuscript

## Data Availability Statement

Original data are available on request

